# Evidence for a long-term protection of wheel-running exercise against cocaine psychomotor sensitization in adolescent but not in young adult mice

**DOI:** 10.1101/188466

**Authors:** Louis-Ferdinand Lespine, Ezio Tirelli

**Author notes:** Corresponding author: Louis-Ferdinand LESPINE, Department of Psychology, University of Liège, Quartier Agora - Place des Orateurs, 2 (B32), 4000 Liège, Belgium (L.F.L).

## Abstract

Rodents housed with a running wheel can exhibit attenuated cocaine seeking and cocaine-induced psychomotor activation. However, the longevity of the exercise anti-drug protection and the influence of the developmental stage during which exercise is displayed received little attention. Here, females and males C57BL/6J mice, aged 28 (adolescents) or 77 (young adults) days were housed with (n=56) or without (n=28) a running wheel. After 3 weeks in these conditions, half of the exercised mice were deprived of their wheel (n=28) whereas the other half and the sedentary mice (no wheel) were kept in their respective environments throughout experimentation. After 3 additional weeks, mice were tested for initiation of psychomotor sensitization to 9 once-daily intraperitoneal injections of 8 mg/kg cocaine (following 2 drug-free test sessions). The expression of sensitization was assessed on a single test session 30 days after the last sensitizing cocaine injection. Continuously exercised mice (wheel throughout experimentation) were less responsive to the initiation and the expression of cocaine effects, regardless of the gender and the developmental period during which exercise was introduced. Wheel-running during adolescence attenuated in later life the initiation and the expression of sensitization in females and only its expression in males. In adult females and males, previously-exercised and sedentary mice exhibited indiscernible levels of initiation and expression of sensitization. Thus, the likelihood of the long-term protection of exercise against cocaine vulnerability may depend not only on the gender but also and especially on the period of life in which exercise took place.

## 1. Introduction

Cross-sectional, clinical and longitudinal studies suggest that regular physical activity could exert preventive effects on the initiation of drugs consumption (Audrain-McGovern et al. 2015; Lisha and Sussman, 2010; Terry-McElrath and O’Malley, 2011) although a causal relationship has yet to be convincingly demonstrated in humans. Animal research reported consistent evidence for beneficial effects of exercise on the vulnerability to several drugs in rats or mice housed for several weeks with a running wheel, the most popular animal model of aerobic exercise. Many authors housed their animals with a continuously-accessible running-wheel throughout experimentation, the assessment of drug responsiveness taking place a number of weeks after the beginning of exercise regime and often outside the home-cage (Bardo and Compton, 2015; Lynch et al. 2013 for reviews). For example, that design was used in studies having reported reduced levels of acquisition or relapse of cocaine or methamphetamine self-administration and of psychomotor sensitization to cocaine (Engelmann et al. 2014; Diaz et al. 2013; Geuzaine and Tirelli, 2014; Lespine and Tirelli, 2015; Smith et al. 2008). Other studies applied the exercise regime entirely before animals were tested for their drug responsiveness. With that procedure, clear-cut attenuating drug effects of a regime of exercise (including daily treadmill activity) have also been reported for self-administration, conditioned place preference, sedation, or stress-induced reinstatement of self-administration induced by amphetamine, ecstasy, morphine, cocaine, nicotine or ethanol (Beiter et al. 2016; Chen et al. 2008; Fontes-Ribeiro et al. 2011; Leasure and Nixon, 2010; Lett et al. 2002; Mollenauer et al. 1991; Ogbonmwan et al. 2015; Rozenske et al. 2011; Sanchez et al. 2015). In these studies, the interval between the end of the exercise regime and the onset of the assessment of drug responsiveness was never longer than a week. Recently, we have founded that a 5-week period of wheel-running in C57BL/6J mice reduced the levels of cocaine psychomotor sensitization initiated 5 days later, suggesting that the anti-drug consequences of exercise don’t vanish immediately following its cessation (Lespine and Tirelli, 2015). Curiously, the longevity of the post-exercise, long-term, protective effects has received little or no attention in spite of its obvious practical importance. Additionally, whatever the timing and duration during experimentation of exercise regime it was often introduced during adolescence or early adolescence, sometimes commencing a few days after weaning (21-25 days of age), and much less frequently during adulthood (in principle after 60 days of age). In fact, it is well recognized that animals may be especially sensitive to internal and external environmental conditions not only of the rearing pre-weaning but also of the post-weaning environment (adolescence and ontogenesis of sexual behavior), exerting a strong and lasting influence on brain and behaviour development (Doremus-Fitzwater et al. 2010; Wahlstrom et al. 2010 for reviews). This is well supported by studies having shown increased neuroplastic processes (such as neurogenesis and angiogenesis) and learning performance in adult rodents that had been exposed as weanlings to free exercise (see review Berchtold et al. 2010; Gomez-Pinilla and Hillman 2013; Merkley et al. 2014). Its seems also that exposing adults to exercise yields smaller short-term and long-term effects than in juveniles and that more intense regimes of exercise are necessary for clear-cut beneficial effects to be induced in older animals. For example, robust improvements of learning-and-memory performance and emotional states have been found in rodents 4 weeks after a 4/5-week regime of wheel-running during adolescence, whereas the same regime of exercise during adulthood did not induce these long-term effects (Hopkins et al. 2011; Jha et al. 2016). Thus, it can be hypothesized that exercise could well be efficacious at inducing long-term anti-drug effects especially when allowed early in life or during adolescence but not, or much less, during adulthood. Here, adolescent and young adult C57BL/6J mice, females and males, were housed with a running wheel for 3 weeks and, 3 weeks later, tested separately for their propensity to exhibit sensitization to the psychomotor-activating effects of cocaine, a phenomenon which likely has an integral role in the mechanisms of craving and relapse (Leyton and Vezina, 2013; Steketee and Kalivas, 2011).

## 2. Experimental procedures

### 2.1. Subjects and basic housing

We employed a total of 336 experimentally-naïve C57BL/6J mice, that were obtained from JANVIER, Le-Genest-Saint-Isle France, either at 21 days of age (just weaned) for Experiments 1 (females, N=84) and 2 (males, N=84) or at 70 days of age for Experiments 3 (females, N=84) and 4 (males, N=84). The choice of C57BL/6J strain was based on its wide use in addiction research and previous experiments performed in our lab. Upon arrival, mice were housed in groups of six in large transparent polycarbonate cages (38.2 x 22 cm surface x 15 cm height; TECHNIPLAST, Milano, Italy) for a period of one week of acclimation. On the following day, mice were housed individually according to the experimental housing conditions (see below Experimental housing conditions section 2.4) in smaller TECHNIPLAST transparent polycarbonate cages (32.5 x 17 cm surface x 14 cm height) with pine sawdust bedding, between-animal visual, olfactory and acoustic interactions remaining possible. Tap water and food (standard pellets, CARFIL QUALITY, Oud-Turnhout, Belgium) were continuously available. The animal room was maintained on a 12:12 h dark-light cycle (lights on at 07.00 a.m.), at an ambient temperature of 20-24°C. In Experiments 1 (females) and 2 (males), mice were aged 71 days and their weight averaged 19.8 g (± 0.16 g standard error of the mean (SEM)) and 22.9 g (± 0.17 g SEM) at the beginning of behavioral testing. In Experiments 3 (females) and 4 (males), mice were aged 120 days and their weight averaged 22.7 g (± 0.18 g SEM) and 27.9 g (± 0.17 g SEM) at the beginning of testing. All experimental treatments and animal maintenance were reviewed by the University of Liège Animal Care and Experimentation Committee (animal subjects review board), which gave its approval according to the Belgian implementation of the animal welfare guidelines laid down by the European Union (“Arrêté Royal relatif à la protection des animaux d’expérience” released on 23 May 2013, and “Directive 2010/63/EU of the European Parliament and of the Council of 22 September 2010 on the protection of animals used for scientific purposes”). All efforts were made to minimize the number of animals used and their suffering. Moreover, the ARRIVE guidelines (Animal Research Reporting In Vivo Experiments), which have been developed to improve the quality of experimenting and reporting in animals studies, were followed as closely as possible (Kilkenny et al. 2010).

### 2.2. Drug treatments

(–)-Cocaine hydrochloride (BELGOPIA, Louvain-La-Neuve, Belgium), dissolved in an isotonic saline solution (0.9% NaCl), was injected at a dose of 8 mg/kg in a volume of 0.01 ml/g of body weight, the control treatment consisting of an equal volume of isotonic saline solution. All injections were given intraperitoneally (i.p.). The dose and route of administration were selected on the basis of our previous study (Lespine and Tirelli, 2015). Note that this dose is known to also induce hedonistic-rewarding effects in mice (Brabant et al. 2005).

### 2.3. Behavioral test chambers

A battery of six home-made chambers, connected to a custom written software for data collection, was used to measure mice psychomotor activity, one mouse being tested in each chamber. Each activity chamber was constituted of a removable transparent polycarbonate tub (22 x 12 cm surface x 12 cm height), embedded onto a black-paint wooden plank serving as a stable base. The lid was made of a transparent perforated acryl-glass tablet. Two photocell sources and detectors were mounted on the plank such that infrared light-beams were located on the two long sides of the tub at 2-cm heights from the floor, 8-cm apart and spaced 6.5 cm from each end of the tub. Psychomotor activity (locomotion) was measured in terms of crossings detected by the beams, one crossing count being recorded every time an ambulating mouse broke successively the two parallel beams. The activity chambers were individually encased in sound-attenuated shells that were artificially ventilated and illuminated by a white light bulb during testing. Each shell door comprised a one-way window allowing periodic surveillance during testing.

### 2.4. Experimental housing conditions

The presence or the absence of a running wheel in each individual home-cage for a given period of time defined the experimental housing conditions, as detailed below in the Experimental design and procedure section 2.5. A running wheel was made of an orange polycarbonate saucer-shaped disk (diameter 15 cm, circumference 37.8 cm; allowing an open running surface) mounted on a plastic cup-shaped base (height 4.5 cm) via a bearing pin so as to being inclined from the vertical plane at an angle of 35° (ENV-044, Med Associates; St Albans, VT, USA). The base was fixed on a stable transparent acryl-glass plate. Running was monitored and recorded once-weekly (for 24 h at an interval of 6 days) by means of a wireless system, each wheel being connected to a USB interface hub (DIG-804, Med Associates) which relayed data to a Wheel Manager Software (SOF-860, Med Associates). To ensure that the amount of physical exercise in the sedentary mice was maintained to a minimum, no locked wheel was placed in their home-cage on which a mouse can display much climbing (Harri et al. 1999).

### 2.5. Experimental design and procedure

Each experiment lasted 85 days, with 43 days devoted to exercise manipulations and the following 42 days to pre-testing (habituation), testing and inter-testing phases (see Fig. 1). Fifty-six mice were initially housed in the aforementioned housing conditions for 3 weeks during adolescence from 28 to 50 days of age (Experiments 1 and 2) or during adulthood from 77 to 99 days of age (Experiments 3 and 4). At the end of three weeks of such a housing, running wheels were removed from half of the home-cages (n=28), in which mice underwent a definitive cessation of exercise. The remaining exercising mice (n=28) kept their wheel until the end of experimentation. Twenty-eight other mice were housed without wheel throughout experimentation, constituting the third experimental housing condition. Note that a group of mice housed for the 3 weeks preceding the beginning of testing, from 50 to 71 days of age (Experiments 1 and 2) or from 99 to 120 days of age (Experiments 3 and 4) was not included for the following reasons. First, the primary purpose and the hypotheses of our experiment did not require it. Second, and more importantly, such a group would have introduced a confounding developmental factor as wheel-running exercise would have taken place in different periods of life, impeding a sound and correct interpretation of the results. Since mice from the three housing environments received cocaine or saline during testing, a basic 3 (exercise) x 2 (cocaine) factorial design was generated for each experiment with n=14. This sample size was determined on the basis of our previous work in which large-effect size interactions between exercise and cocaine were found for acute responsiveness and sensitization (initiation and expression) to the psychomotor-stimulating effect of the drug (minimum effect sizes at ^2^p = .11 or *d* = 0.70; recalculated from Lespine and Tirelli, 2015). Each mouse was assigned to one of the six experimental groups by means of a computer-generated randomization schedule, the six mice housed in the acclimation cages contributing to the six possible treatments in as many different experimental groups defined as follows. Group SED/COC: mice living without wheel throughout experimentation and receiving cocaine during testing; group SED/SAL: mice living without wheel throughout experimentation and receiving saline during testing; group TEX/COC: mice exercising throughout experimentation receiving cocaine during testing; group TEX/SAL: mice exercising throughout experimentation receiving saline during testing; group 3EX/COC: mice exercising only during the first three weeks of experimentation and receiving cocaine during testing; group 3EX/SAL: mice exercising only during the three first weeks of experimentation and receiving saline during testing. To take into account any between-session variability due to the time and circumstances of testing, the six experimental conditions were systematically represented within each test session by one mouse. In other words, each test session formed a block including six mice individually tested in as many activity-chambers. Since there were 14 mice per group in all experiments, each experiment involved 14 blocks that were incorporated into a randomized block design. Each experiment was divided in two identical sub-experiments involving seven daily sessions (blocks) performed successively between 08.30 a.m. and 01.00 p.m. After each test session, animals were returned to the animal room within 10 min and the activity-chambers were cleaned with a disinfectant. One week before the beginning of testing, all mice were injected once with a saline solution in the animal room to familiarize them with that manipulation. Behavioral testing included the following four phases articulated over a 42-day period. (1) A pre-test basal session to familiarize animals to the novelty of the test context without neither injection nor measures. (2) A two-session test assessing the psychomotor-activating effects of cocaine measured under saline (2^nd^ once-daily session) and after a first injection of cocaine (3^rd^ once-daily session). (3) Eight once-daily injections of cocaine, with the measurement of the psychomotor-activating effect of the drug after each injection, initiating psychomotor sensitization after the 3^rd^ session (total of 9 sessions). (4) Taking place 30 days after the 9^th^ (last) cocaine injection, a unique test assessing the long-term expression of sensitization on which animals received their previous respective pharmacological treatment. This test took place on the last day of the experimentation period (see Fig.1 for a schematic representation of the experiment time-line). Throughout testing, mice were weighted and received their pharmacological treatment right before being placed in the test chamber, the recording of ambulatory crossings lasting 30 min in all sessions. Experimental blinding was not realized because the unique experimenter inevitably knew the housing condition of each mouse and whether cocaine or saline has to be administered. We believed that a complicated coding with a unique experimenter would not have substantially improved the quality of testing. Moreover, psychomotor activity was recorded automatically, minimizing the possibility of a direct influence of the experimenter on the data collection.

**Figure 1.**
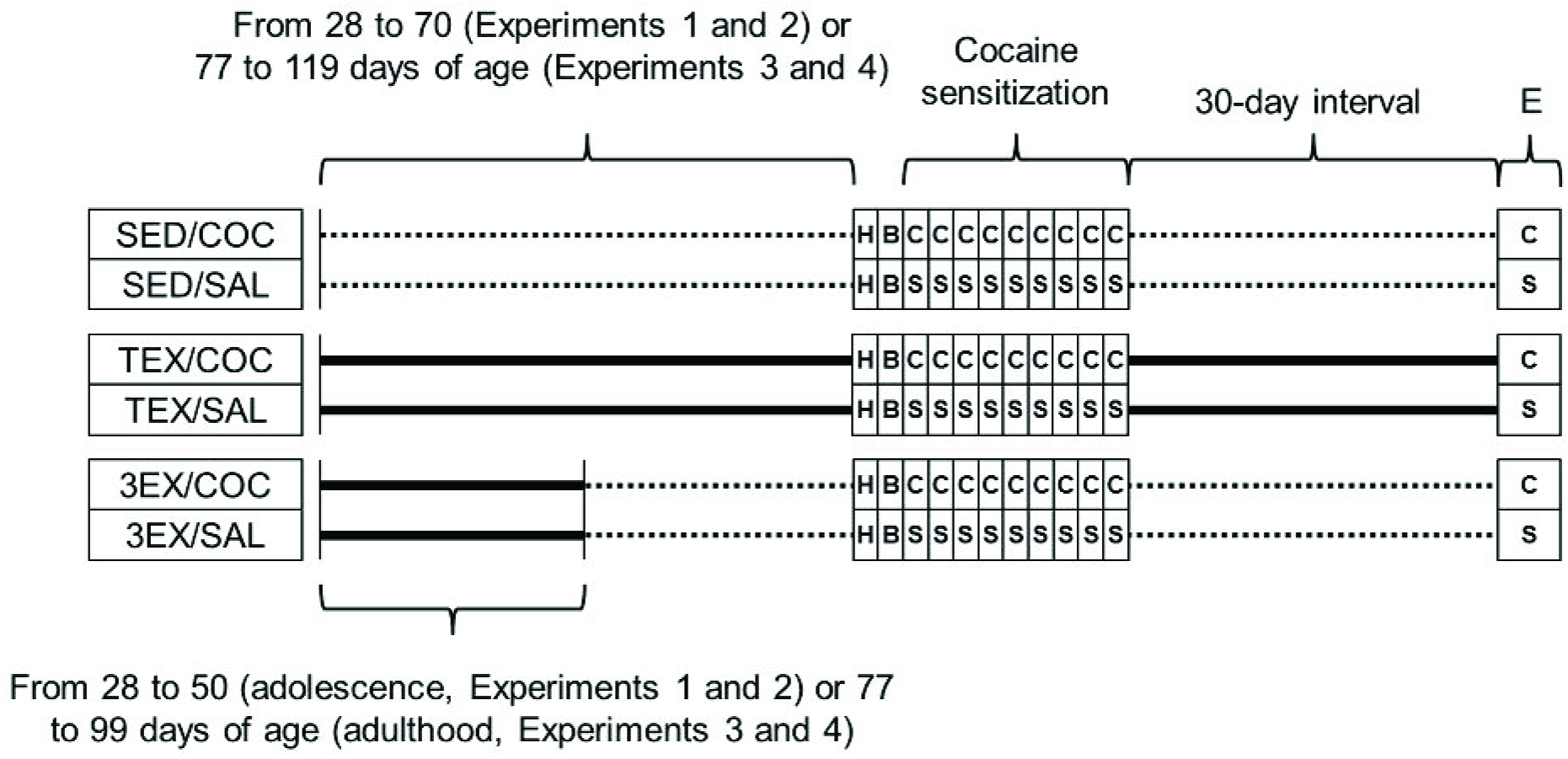
Experimental timeline and design. Mice were housed individually either in the presence (n=56) or the absence (n=28) of a running wheel for at least 3 weeks from the 28^th^ to the 50^th^ day of age (Experiments 1 and 2) or from the 77^th^ to the 99^th^ day of age (Experiments 3 and 4). The day following this period, half (n=28) of the mice housed with a running wheel were switched to housing conditions without wheel (3EX/COC and 3EX/SAL groups; n=14), the other half of the exercised mice (n=28) remaining with their wheel until the end of experimentation (TEX/COC and TEX/SAL groups; n=14). Sedentary, unexercised, mice (n=28) lived without running wheel throughout experimentation (SED/COC and SED/SAL groups; n=14). Testing began after an interval of 3 additional weeks. Thus, each experiment comprised six experimental groups with N=84, mice from each housing group receiving either cocaine or saline. Solid lines represent the presence of a running wheel in the home-cage and dotted lines its absence. H: habituation session (to familiarize animals to the novelty of the test context without neither injection nor measures); B: baseline session; the 2^nd^ once-daily session assessing the baseline activity under saline; C: cocaine intraperitoneal administration (repeated 8 times, once-daily); S: saline administration (repeated 8 times, once-daily). E: single-session test on which the long-term expression of the sensitization was assess 30 days after the last sensitizing injection, under the previous pharmacological treatments.

### 2.6. Data analysis

Inferential statistics were computed on the following data: (1) the acute responsiveness to the psychomotor-activating effects of cocaine scored as the absolute difference between the values (number of crossings) derived from the first cocaine session and those of the baseline session under saline, (2) the overall responsiveness to the psychomotor-activating effects of cocaine over the first session and the additional 8 injections (initiation of sensitization) scored as the area under the curve with respect to zero (AUC ground; calculation formula detailed in Pruessner et al. 2003). (3) The psychomotor activity (scored as the number of crossings) exhibited during the (last) test session revealing the expression of sensitization. Data dealing with the initiation (development) of sensitization are only presented in the figures and were not statistically analysed. Each set of data was analysed according to a randomized block design with a fixed-model 3x2 ANOVA incorporating the housing condition (period of exercise; 3 levels) and pharmacological treatment (cocaine or saline; 2 levels) as between-group factors, with the session as a blocking factor (14 levels since n = 14). The primary outcomes, defined by our focused hypotheses (described in the Experimental design and procedure section 2.5), were given by a limited number of planned crossed contrasts that contributed to the interaction (see Rosenthal and Rosnow, 2009). Specifically, our crossed contrasts compared the difference (effect) between the values of the TEX/COC and TEX/SAL groups or those of the 3EX/COC and 3EX/SAL groups with the difference (effect) between the values of the SED/COC and SED/SAL groups, each contrast being derived from the mean-square error term (MSE) provided by the ANOVA. It is on these bases (differences between these two effects or between-group differences) that effect sizes were estimated and given by Cohen’s *d* (calculated from the corresponding *t*s and *df*s). Note that magnitudes of the Cohen’s *d* are conventionally classified as very small (0.01), small (0.20), medium (0.50), large (0.80), very large (1.2) and huge (2.0) (see Sawilowsky, 2009). One-tailed *t*-tests were used for these comparisons since SED/COC mice were expected to display greater cocaine responsiveness than exercised mice, statistical significance being set at *p* = .025 (Bonferroni correction). As secondary outcomes, the basic psychomotor-activating effects of cocaine within each housing (exercise) condition was ascertain using one-tailed *t*-tests taken at *p* = .05, which compared the values derived from the TEX/COC, 3EX/COC and SED/COC groups with those of their respective control TEX/SAL, 3EX/SAL and SED/SAL groups (*p*-values adjustments were unnecessary since such effects are usually oversized). For the long-term expression of sensitization, representing the final experimental outcome, the degree of precision associated with the effect size estimates was given by 95% confidence intervals (CI) derived from non-centrality interval estimation for the non-central *t* distributions (Steiger, 2004). Except for the effect size estimates, means ± SEM are presented in the graphs.

## 3. Results

### 3.1. Wheel-running activity prior to behavioral testing

As shown in Fig. 2, all mice housed with a wheel (TEX and 3EX) showed a rapid increase in wheel-running (expressed in kilometres) over the two first weeks until reaching a plateau. At this stage of the experiment, mice from the cocaine and saline groups (within each housing condition) were still undistinguishable and pooled such that n=28. Mice continuously housed with a wheel exhibited a slight decrease as from the 36^th^ day, leading to an inverted-U shape, a pattern which has been already reported in C57BL/6J (Fuss et al. 2010). Note that previous observations in our laboratory indicate that mice tend to plateauing rather than decreasing their running activity (unpublished results). The pattern was comparable across all the groups and appears similar to that reported in our prior study (Lespine and Tirelli, 2015) and other unpublished data from our lab. As expected, there were no discernible between-group differences related to the duration of wheel-running exercise, the use of inferential statistics being unnecessary.

**Figure 2.**
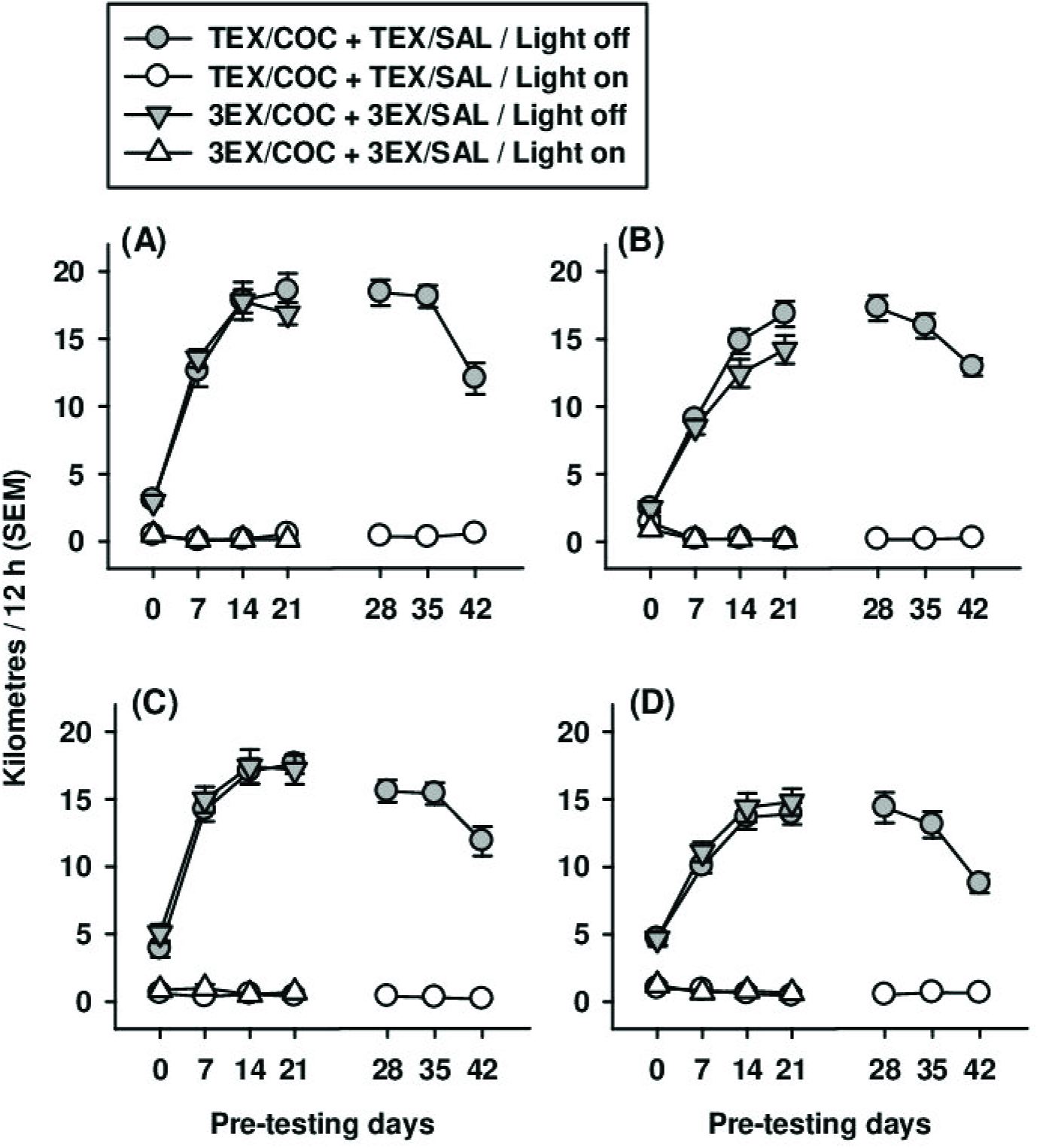
Wheel-running activity recorded prior to the testing period. Nocturnal (light off) and diurnal (light on) wheel-running activity during the pre-testing period for the two possible regimes of exercise: 3 weeks of (discontinuous) exercise (3EX/COC and 3EX/SAL groups, females or males) or continuous exercise (TEX/COC and TEX/SAL groups, females or males). (A) Distances travelled by females for whom a running wheel was introduced at 28 days old (experiment 1). (B) Distances travelled by males for whom a running wheel was introduced at 28 days old (experiment 2). (C) Distances travelled by females for whom a running wheel was introduced at 77 days (experiment 3). (D) Distances travelled by males for whom a running wheel was introduced at 77 days (experiment 4). The horizontal axis indicates the last day of each of the 6 weeks of housing with a running wheel (7^th^, 14^th^, 21^st^, 28^th^, 35^th^ and 42^nd^ days; 24h each), the day denoted by 0 being the starting day preceding the 3-week (3EX groups) or 6-week (TEX groups) pretesting period. No inferential statistics were conducted on these data.

### 3.2. Experiment 1. Exercise introduced at 28 days of age (adolescence) in females

Panel A in Fig. 3 depicts the results dealing with the initiation of cocaine psychomotor sensitization over 9 once-daily sessions in females housed either with (TEX/COC, TEX/SAL, 3EX/COC and 3EX/SAL groups) or without (SED/COC and SED/SAL groups) a running wheel during adolescence (as from 28 days of age) and tested 6 weeks later (at 71 days of age), 3EX/COC and 3EX/SAL mice having stopped wheel-running exercise 3 weeks before testing (at 50 days of age). These data were not statistically analysed. Panel B presents the acute responsiveness assessed on the 3^rd^ day of exposure of the mice to the activity-chambers. Crossed contrasts suggested that the psychomotor-activating effect of cocaine was robustly (relatively large effect sizes) attenuated in both continuously (TEX/COC vs. TEX/SAL groups) and discontinuously (3EX/COC vs. 3EX/SAL groups) exercised mice in comparison with the effect found in the sedentary or unexercised (SED/COC vs. SED/SAL groups) animals (*d* = 0.71, *t*(65) = 2.88, *p* = .003 and *d* = 0.95, *t*(65) = 3.84, *p* < .001, respectively). Panel C depicts the overall responsiveness during the initiation phase of cocaine sensitization (9-session development) scored as AUC ground. The pattern of interactive effects was comparable to that found for the acute psychomotor-activating effect of cocaine, but with greater effect sizes (*d* = 0.96, *t*(65) = 3.85 and *d* = 1.19, *t*(65) = 4.78 both at *p* < .001, respectively). Panels D and E depict the results on the single-session test for the expression of sensitization and the related effect sizes (crossed contrasts). The psychomotor responses to cocaine were substantially smaller among the exercised cocaine-sensitized mice (TEX/COC vs. TEX/SAL or 3EX/COC vs. 3EX/SAL groups) than in their sedentary counterparts (SED/COC vs. SED/SAL groups). The interaction-related differences between the effects obtained in the exercised groups and that found in the sedentary animals were both very large since *d* = 1.62 with 95% CI 1.06-2.18 (*t*(65) = 6.55, *p* < .001) and *d* = 1.42 with 95% CI 0.87-1.96 (*t*(65) = 5.72, *p* < .001). Statistics related to the three basic psychomotor-activating effects of cocaine within each housing (exercise) condition (TEX/COC vs. TEX/SAL, 3EX/COC vs. 3EX/SAL and SED/COC vs. SED/SAL groups) are provided in Supplement 1, Table 1 (secondary outcomes).

**Figure 3.**
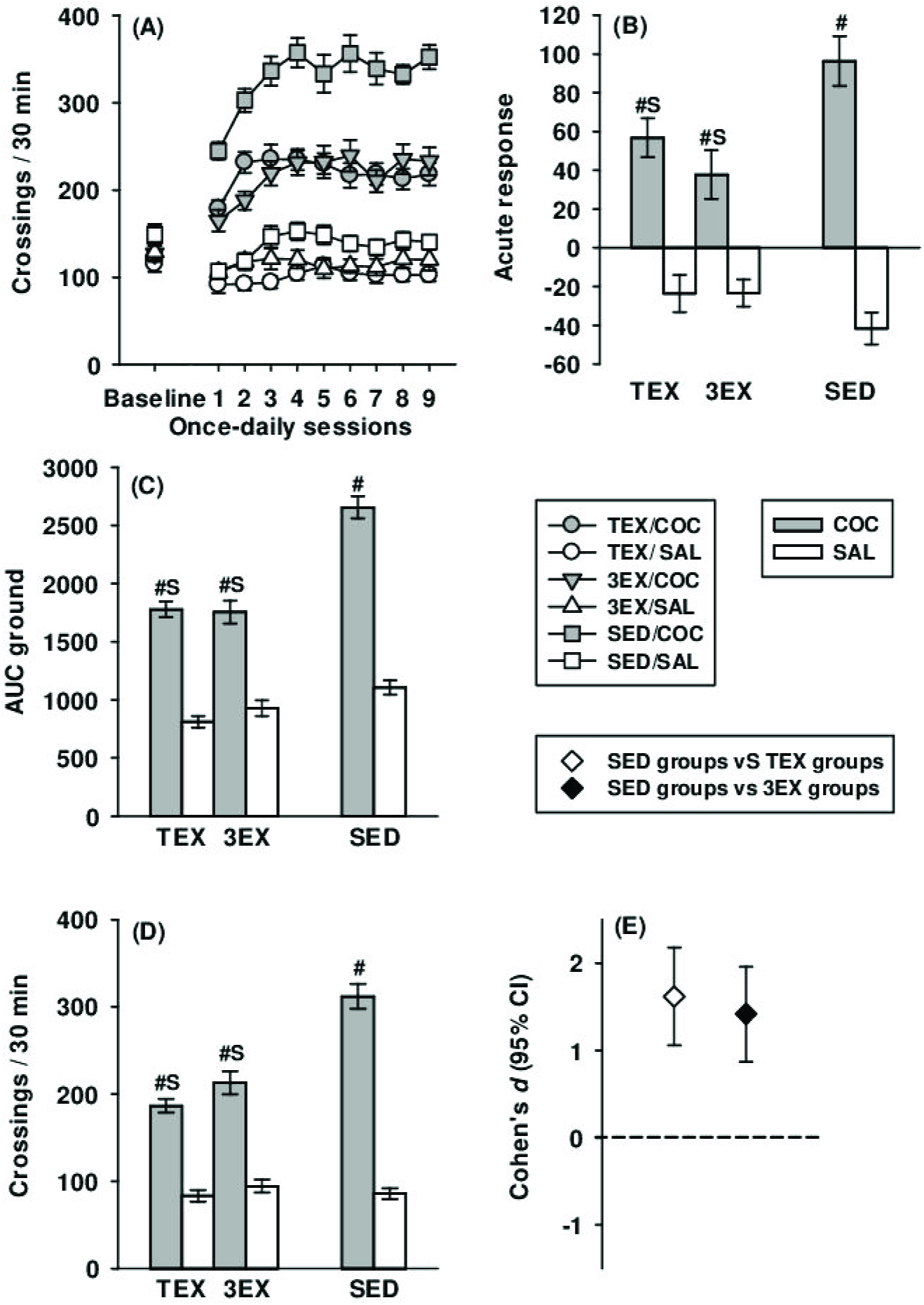
Experiment 1: cocaine psychomotor responsiveness in female mice housed with a running wheel for 3 weeks during adolescence (as from the 28 days of age) and tested at 71 days of age. (A) Acute psychomotor responsiveness to cocaine scored as the absolute difference between values from the first cocaine session and the baseline session. (B) Baseline psychomotor activity (under saline) and initiation (development) of psychomotor sensitization over 9 once-daily sessions (the data from the acute responsiveness being those plotted for the first once-daily cocaine injection). (C) Cocaine-induced psychomotor levels measured over the initiation (development) of sensitization (overall responsiveness) scored as AUC ground. (D) The increment of the psychomotor levels over the cocaine repeated injections scored as AUC increase. (E) Psychomotor responsiveness on the single-session test for long-term expression of cocaine sensitization. (F) Effect sizes (Cohen’s *d* with 95% CI) corresponding to the crossed contrasts assessing the interactions between cocaine and exercise in the results dealing with the long-term expression of cocaine sensitization (the effect sizes associated to the other measures of cocaine responsiveness are indicated in the text). (S): significant interaction-related difference between the effects displayed by the exercised mice (TEX/COC vs. TEX/SAL and 3EX/COC vs. 3EX/SAL groups) and that measured in the sedentary mice (SED/COC vs. SED/SAL) taken at a *p*-level of .025. (#): significant difference between the two groups having received cocaine or saline within each exercise condition taken at a *p*-level of .05.

**Table 1.** Summary of the results regarding the post-exercise long-term (interval of 3 weeks) attenuating effects of a 3-week wheel-running regime on cocaine psychomotor responsiveness and sensitization. **YES**: cocaine effect in the exercised mice (3EX/COC vs 3EX/SAL) detected as significantly smaller (with interaction crossed contrasts) than that displayed by the sedentary mice (SED/COC vs SED/SAL). **NO**: no detection of a significant difference between these two cocaine effects (no interaction between exercise and cocaine). The corresponding Cohen’s *d* effect size is indicated in parentheses.

|  | Acute (first<br>response | Initiation | Expression |
| --- | --- | --- | --- |
| Exp. 1. Exercise introduced at 28 days of<br>age (adolescence) in females | **YES** (0.95) | **YES** (1.19) | **YES** (1.42) |
| Exp. 2. Exercise introduced at 28 days of<br>age (adolescence) in males | NO (0.42) | NO (0.28) | **YES** (0.53) |
| Exp. 3. Exercise introduced at 77 days of<br>age (adulthood) in females | NO (0.31) | NO (-0.12) | NO (0.10) |
| Exp. 4. Exercise introduced at 77 days of<br>age (adulthood) in males | NO (-0.10) | NO (0.03) | NO (-0.08) |

### 3.3. Experiment 2. Exercise introduced at 28 days of age (adolescence) in males

Panel A in Fig. 4 shows the results dealing with the initiation of cocaine psychomotor sensitization over 9 once-daily sessions in males living either with (TEX/COC, TEX/SAL, 3EX/COC and 3EX/SAL groups) or without (SED/COC and SED/SAL groups) a running wheel during adolescence (as from 28 days of age) and tested 6 weeks later (at 71 days of age), 3EX/COC and 3EX/SAL mice having stopped wheel-running exercise 3 weeks before testing (at 50 days of age). Panel B depicts the acute psychomotor-activating effects of cocaine (first cocaine injection). These were statistically comparable across the continuously and discontinuously exercised, and sedentary conditions (*t*(65) = 1.81, *p* = .037 and t(65) = 1.69, *p* = .048). The effect sizes of the interaction-related differences between the cocaine effects found in the two exercised groups and that obtained in the sedentary groups can be considered as practically medium (*d* = 0.45 and *d* = 0.42). Panel C shows the overall responsiveness (AUC ground) during the initiation of cocaine sensitization. Crossed contrasts indicate that the continuously exercised mice displayed a significantly smaller responsiveness to cocaine than the sedentary mice (*d* = 0.88, *t*(65) = 3.53, *p* < .001). However, the effect displayed by the mice exercised during 3 weeks did not differ statistically from that found for the sedentary animals, the effect size being small (*d* = 0.28, *t*(65) = 1.11, *p* = .14). Panel D presents the results of the single-session test for the expression of the sensitization to cocaine and Panel E depicts the corresponding effects sizes underlying the interactions (crossed contrasts). That response was significantly smaller in the continuously exercised mice than in the unexercised animals, the effect size being clearly large (*d* = 0.96 with 95% CI 0.45-1.48; *t*(65) = 3.89, *p* < .001). Mice with a history of 3-week exercise also exhibited a significantly lower cocaine expression-related responsiveness, but with a medium effect size (*d* = 0.53 with 95% CI 0.04-1.03; *t*(65) = 2.15, *p* = .018). Statistics related to the three basic psychomotor-activating effects of cocaine within each housing (exercise) condition are provided in Supplement 1, Table 2 (secondary outcomes).

**Figure 4.**
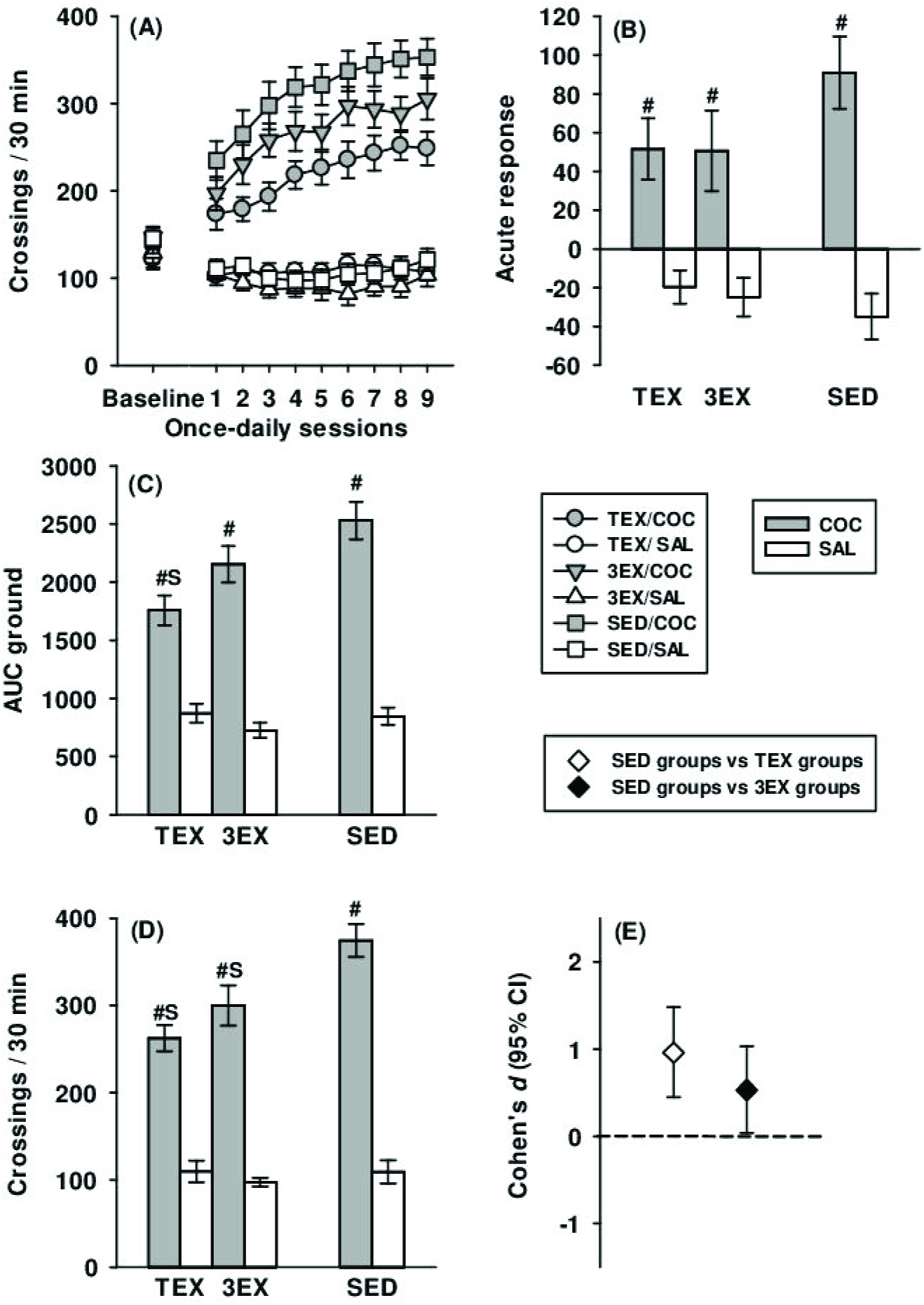
Experiment 2: cocaine psychomotor responsiveness in male mice housed with a running wheel for 3 weeks during adolescence (as from 28 days of age) and tested at 71 days of age. (A) Acute psychomotor responsiveness to cocaine scored as the absolute difference between values from the first cocaine session and the baseline session. (B) Baseline psychomotor activity (under saline) and initiation (development) of psychomotor sensitization over 9 once-daily sessions (the data from the acute responsiveness being those plotted for the first once-daily cocaine injection). (C) Cocaine-induced psychomotor levels measured over the initiation (development) of sensitization (overall responsiveness) scored as AUC ground. (D) The increment of the psychomotor levels over the cocaine repeated injections scored as AUC increase. (E) Psychomotor responsiveness on the single-session test for long-term expression of cocaine sensitization. (F) Effect sizes (Cohen’s *d* with 95% CI) corresponding to the crossed contrasts assessing the interactions between cocaine and exercise in the results dealing with the long-term expression of cocaine sensitization (the effect sizes associated to the other measures of cocaine responsiveness are indicated in the text. (S): significant interaction-related difference between the effects displayed by the exercised mice (TEX/COC vs. TEX/SAL and 3EX/COC vs. 3EX/SAL groups) and that measured in the sedentary mice (SED/COC vs. SED/SAL) taken at a *p*-level of .025. (#): significant difference between the groups having received cocaine or saline within each exercise condition taken at a *p*-level of .05.

### 3.4. Experiment 3. Exercise introduced at 77 days of age (adulthood) in females

Panel A in Fig. 5 depicts results dealing with the initiation of cocaine psychomotor sensitization over 9 once-daily sessions in females living either with (TEX/COC, TEX/SAL, 3EX/COC and 3EX/SAL groups) or without (SED/COC and SED/SAL groups) a running wheel during adulthood (as from 77 days of age) and tested 6 weeks later (at 120 days of age), 3EX/COC and 3EX/SAL mice having stopped wheel-running exercise 3 weeks before testing (at 99 days of age). As shown in Panel B, no significant interaction-related differences between the acute psychomotor effects exhibited by the exercised mice (TEX or 3EX groups) and that found for the sedentary mice (SED groups), were detected by the crossed contrasts, the two effect sizes being small to medium (*d* = 0.38, *t*(65) = 1.52, *p* = .07 and *d* = 0.31, *t*(65) = 1.26, *p* = .11). The results dealing with the overall responsiveness to cocaine during the initiation of sensitization (AUC ground) are graphed in Panel C. As compared with the cocaine effect found in the sedentary mice, continuously but not discontinuously exercised (3-week regime of wheel-running) mice displayed a medium and significantly reduced overall cocaine responsiveness (*d* = 0.52, *t*(65) = 2.09, *p* = .02; *d* = −0.12, *t*(65) = −0.48, *p* = .32). Panels D show the results obtained in the single-session test for the expression of sensitization and Panel E presents the corresponding effect sizes underlying the interactions (crossed contrasts). That cocaine response was significantly lower in the continuously exercised than in the sedentary animals, the corresponding effect size being patently large (*d* = 1.02, 95% CI 0.50-1.53, *t*(65) = 4.11, *p* < .001). By contrast, the difference between the effect found in mice with a history of 3 weeks of exercise and that exhibited by the sedentary mice failed to achieve statistical significance and the corresponding effect size was small (*d* = 0.10, 95% CI −0.39-0.58, *t*(65) = 0.39, *p* = .35). Statistics related to the three basic psychomotor-activating effects of cocaine within each housing (exercise) condition are provided in Supplement 1, Table 3 (secondary outcomes).

**Figure 5.**
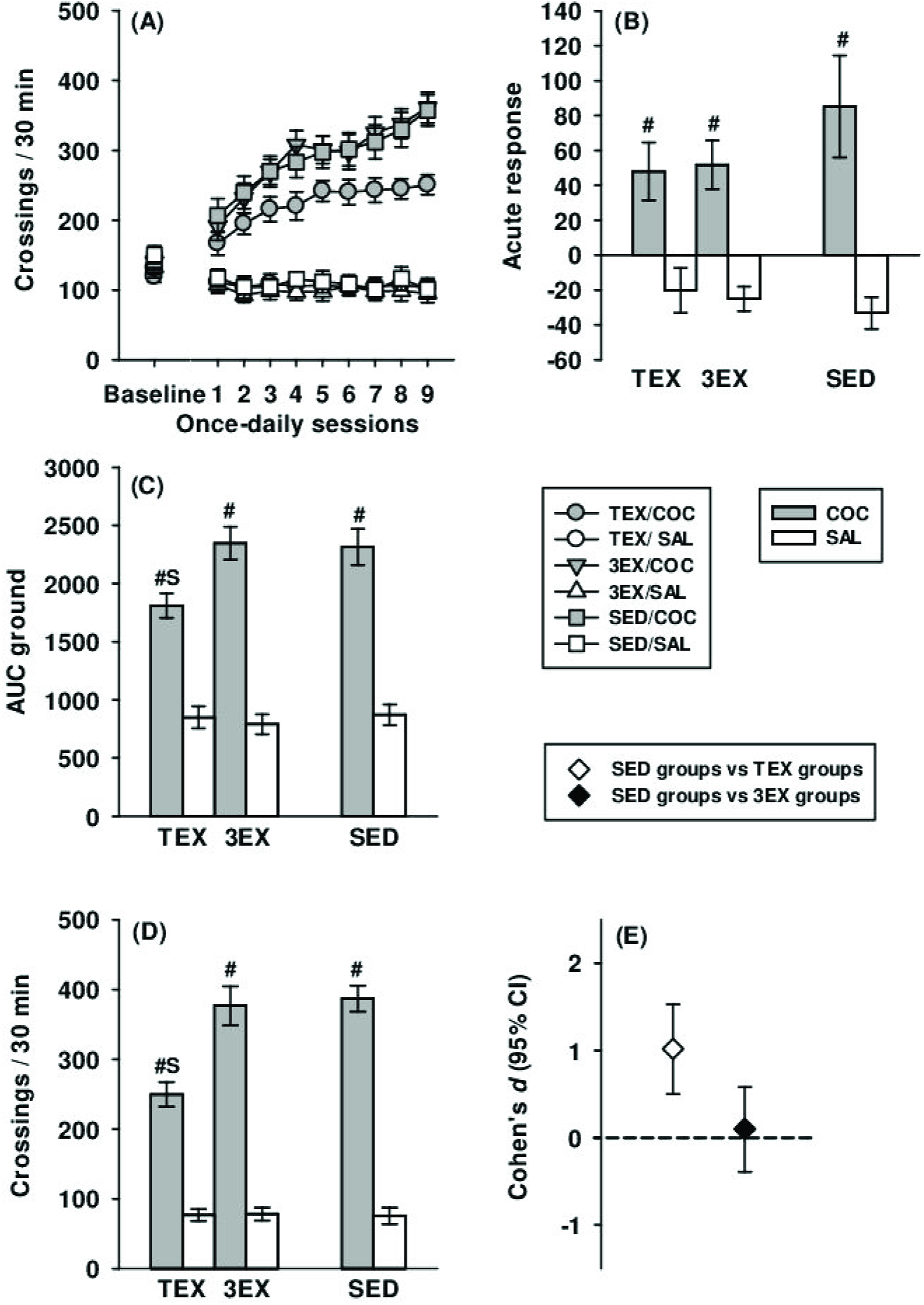
Experiment 3: cocaine psychomotor responsiveness in female mice housed with a running wheel during adulthood (as from 77 days of age) and tested at 120 days of age. (A): Acute psychomotor responsiveness to cocaine scored as the absolute difference between values from the first cocaine session and the baseline session. (B): Baseline psychomotor activity (under saline) and initiation (development) of psychomotor sensitization over 9 once-daily sessions (the data from the acute responsiveness being those plotted for the first once-daily cocaine injection). (C) Cocaine-induced psychomotor levels measured over the initiation (development) of sensitization (overall responsiveness) scored as AUC ground. (D): The increment of the psychomotor levels over the cocaine repeated injections scored as AUC increase. (E): Psychomotor responsiveness on the single-session test for long-term expression of cocaine sensitization. (F): Effect sizes (Cohen’s *d* with 95% CI) corresponding to the crossed contrasts assessing the interactions between cocaine and exercise in the results dealing with the long-term expression of cocaine sensitization (the effect sizes associated to the other measures of cocaine responsiveness are indicated in the text). (S): significant interaction-related difference between the effects displayed by the exercised mice (TEX/COC vs. TEX/SAL and 3EX/COC vs. 3EX/SAL groups) and that measured in the sedentary mice (SED/COC vs. SED/SAL) taken at a *p*-level of .025. (#): significant difference between the two groups having received cocaine or saline within each exercise condition taken at a *p*-level of .05.

### 3.5. Experiment 4: Exercise introduced at 77 days of age (adulthood) in males

Panel A in Fig. 6 depicts results dealing with the initiation of cocaine psychomotor sensitization (incremental development) over 9 once-daily sessions in males living either with (TEX/COC, TEX/SAL, 3EX/COC and 3EX/SAL groups) or without (SED/COC and SED/SAL groups) a running wheel during adulthood (as from 77 days of age) and tested 6 weeks later (at 120 days of age), 3EX/COC and 3EX/SAL mice having stopped wheel-running exercise 3 weeks before testing (at 99 days of age). Panel B depicts the results dealing with the acute psychomotor-activating effects of cocaine (first cocaine injection). The continuously exercised mice displayed a significantly smaller acute responsiveness to cocaine than the sedentary mice, that difference being large-sized (*d* = 0.83, *t*(65) = 3.33; *p* <.001). However, the effect exhibited by the discontinuously exercised mice did not differ statistically from that found for the unexercised animals, the effect size being negligible (*d* = −0.10, *t*(65) = −0.42, *p* = .34). As shown in Panel C, a comparable profile of effects was found for the overall responsiveness to cocaine during sensitization (AUC-ground measure). The continuously exercised mice displayed a significantly smaller overall responsiveness to cocaine than the sedentary mice, the difference being large-sized (*d* = 0.79, *t*(65) = 3.19; *p* < .001). However, the effect exhibited by the discontinuously (3 weeks with a running wheel) exercised mice did not differ statistically from that found for the unexercised sedentary animals, the effect size being negligible (*d* = 0.03, *t*(65) = 0.13, *p* = .45). Panel D presents the results of the single-session test for the expression of cocaine sensitization. While a significant difference was detected between the cocaine effects of the continuously exercised and the unexercised sedentary animals (*t*(65) = 2.77, *p* = .004), the cocaine effect shown by the unexercised groups did not differ from that of the mice exercised during 3 weeks, (*t*(65) = −0.34, *p* = .37). Panel F shows that the corresponding interaction-related effects sizes (crossed contrasts) were medium to large for the continuously exercised mice (*d* = 0.69, 95% CI 0.18-1.19) or very small (and negative) for the mice having used the running wheel for 3 weeks (*d* = −0.08, 95% CI −0.57-0.40). Statistics related to the three basic psychomotor-activating effects of cocaine within each housing (exercise) condition are provided in Supplement 1, Table 4 (secondary outcomes).

**Figure 6.**
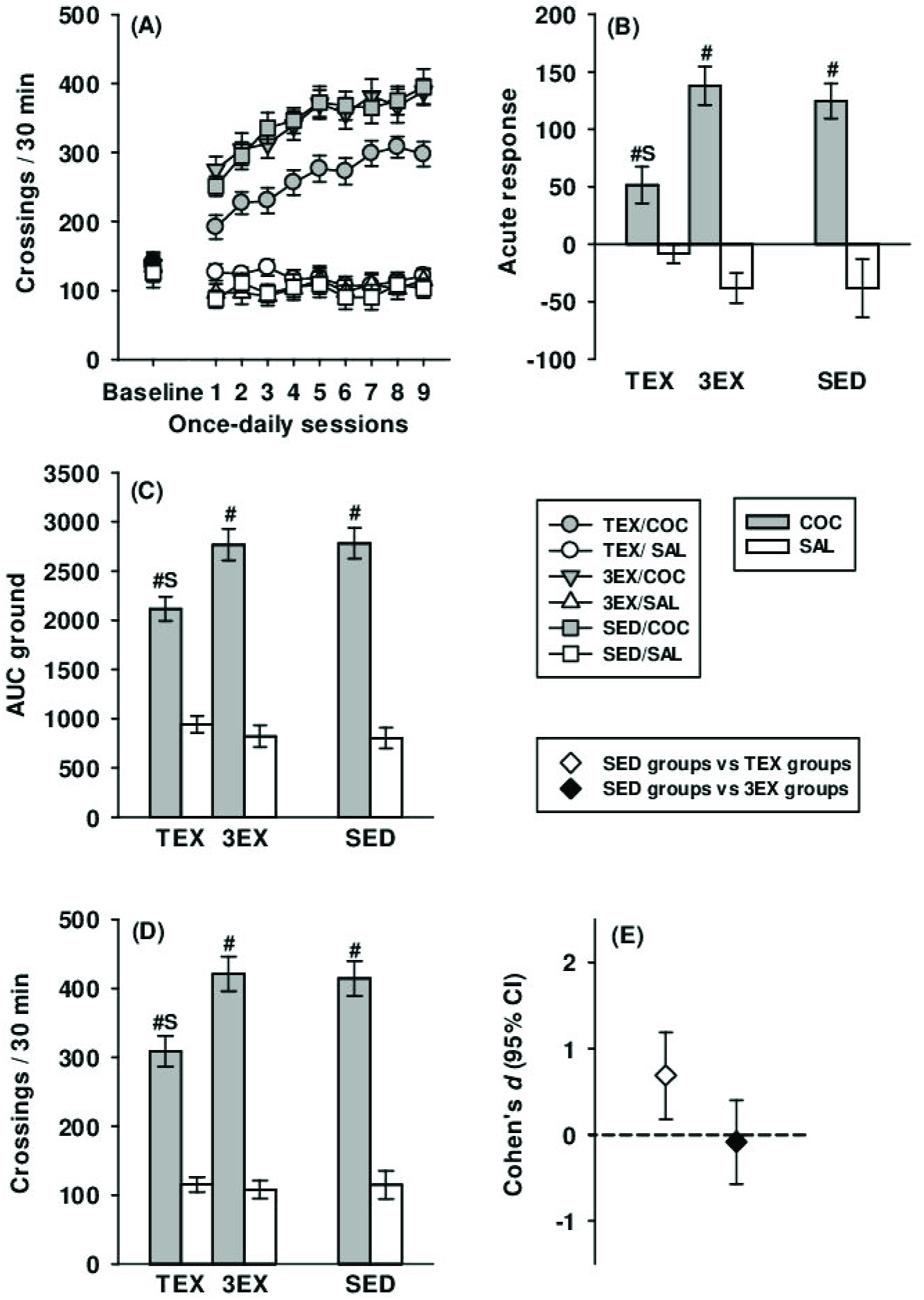
Experiment 4: cocaine psychomotor responsiveness in male mice housed with a running wheel during adulthood (as from 77 days of age) and tested at 120 days of age. (A) Acute psychomotor responsiveness to cocaine scored as the absolute difference between values from the first cocaine session and the baseline session. (B) Baseline psychomotor activity (under saline) and initiation (development) of psychomotor sensitization over 9 once-daily sessions (the data from the acute responsiveness being those plotted for the first once-daily cocaine injection). (C) Cocaine-induced psychomotor levels measured over the initiation (development) of sensitization (overall responsiveness) scored as AUC ground. (D) The increment of the psychomotor levels over the cocaine repeated injections scored as AUC increase. (E) Psychomotor responsiveness on the single-session test for long-term expression of cocaine sensitization. (F) Effect sizes (Cohen’s *d* with 95% CI) corresponding to the crossed contrasts assessing the interactions between cocaine and exercise in the results dealing with the long-term expression of cocaine sensitization (the effect sizes associated to the other measures of cocaine responsiveness are indicated in the text). (S): significant interaction-related difference between the effects displayed by the exercised mice (TEX/COC vs. TEX/SAL or 3EX/COC vs. 3EX/SAL groups) and that measured in the sedentary mice (SED/COC vs. SED/SAL) taken at a *p*-level of .025. (#): significant difference between the two groups having received cocaine or saline within each exercise condition taken at a *p*-level of .05.

## 4. 4. Discussion

Table 1 summarizes the most important findings of the present study. We founded age- and gender-related dissimilarities in the efficacy of 3-week exercise at curbing cocaine sensitization assessed after a waiting period of 3 other weeks (long-term effects). Strikingly, wheel-running was patently efficacious when introduced during adolescence (Cohen’s *d* up to 1.42), but much less so, or more ambiguously, when allowed during early adulthood (Cohen’s *d* ranging from −0.12 to 0.31). In females exercised as adolescents, the 95% CI range of values for the strength of the exercise-by-cocaine interaction ranged from strong to very strong effects (0.87-1.96). In the corresponding males, such protective effects of previous wheel-running were significant only for the expression of sensitization (with a moderate *d* of 0.53). Notice that there were no convincing relationships (very small and non-significant correlations and slopes, not shown) between running distances and any of the three psychopharmacological measures, an outcome in accord with previous results (Lespine and Tirelli, 2015; Geuzaine and Tirelli, 2014; Smith and Pitts, 2011).

Our hypothesis was thus confirmed, these results constituting to our knowledge the first report of a relatively enduring (several weeks), long-term, protection against drug effects induced by a prior period of exercise. That these effects occurred only in mice exercised as adolescents is in substantial accord with non-pharmacological studies reporting that an early regime of exercise can entail long-term positive effects on brain and behaviour, which remained unobserved or barely detectable in animals exercised as adults (aged more than 60 days during the exercise period). For example, in the Hopkins et al. study (2011), adolescent rats housed with a running wheel for 4 weeks (as from 32 days of age) showed an enhanced performance in object recognition memory not only shortly after exercise regime but also 4 weeks later, whereas similarly-exercised (as from 63 days of age) adult rats were unable to do so at the longest post-exercise interval (concomitantly BDNF levels were heightened in the perirhinal cortex, which is indispensable for memory processes). Similarly, adolescent (aged less than 2 months) genetically-depressed mice initially housed for 2 months with a running wheel as part of a composite environment exhibited 4 weeks later reduced scores of depression-like behaviours, whereas their adult counterparts aged 2-4 or 12-14 months did not so (Jha et al. 2016).

As in the psychopharmacology studies showing exercise protective effects (above), rodents used in the neurobehavioral studies that reported long-term improving effects of wheel-running on ulterior endpoints (intervals of 3-5 weeks) received the exercise regime during adolescence, when aged less than 60 days (Berchtold et al. 2010; Gomez-Pinilla and Hillman, 2013 Merkley et al. 2014). Similarly, in a recent study, similarly-aged rats housed in a composite environment with wheels impeded the full development of allodynia induced by chronic constriction injury (a model of neuropathic pain) for up to almost 3 months after the removal of this environment (Grace et al. 2016).

Although the literature on the long-term neurobehavioral and psychopharmacological consequences of exercise early in life is still meagre, one can admit that these are globally compatible with the conception of a great malleability of the brain in maturing mammals. The attenuation in cocaine responsiveness observed in our mice exercised as adolescents (and well-marked in females) may be related to structural and functional brain plasticity in regions involved in motivation and reward (among others) that continue to mature during adolescence (see Doremus-Fitzwater et al. 2010; Gomez-Pinilla and Hillman 2013; Wahlstrom et al. 2010 for reviews). This is of course also true for adverse events experienced early in life, known to entail negative long-term developmental effects. For example, chronic stressful experiences during adolescence are associated with higher risk to develop substance abuse disorders (Jordan and Andersen, 2017).

The post-exercise decreased cocaine effects described here, and in our previous study (Lespine and Tirelli, 2015), strongly contrast with the previously-reported increase in methamphetamine self-administration or cocaine conditioned place preference occurring shortly after the removal of the running-wheel or the housing environment comprising a wheel (Engleman et al. 2014; Nader et al. 2012). In these studies, the post-exercise increase in drug vulnerability was seen as a result of the presumably stressful consequences of the running-wheel deprivation, exercise being conceived as an anti-stress factor (Solinas et al. 2010). This anti-stress view of the anti-drug effects of exercise is partly based on the well documented accentuating effects that classical stressors induce on drug vulnerability (Burke and Miczek, 2013; McCormick et al. 2010) and is also backed by non-pharmacological studies having reported an increase in anxiety after exercise cessation in mice (Greenwood et al. 2012; Malisch et al. 2009; Nishijima et al. 2013). Therefore, our results (including those published in Lespine and Tirelli, 2015), along with several other studies (e.g. Beiter et al. 2016; Mollenauer et al. 1991; Sanchez et al. 2015), do not support the anti-stress view of the anti-drug effects of exercise.

It is reasonable to admit that even a small beneficial (long-term) effect of discontinuous exercise is something gained and noteworthy (and not trivial). However, to be detected as statistically significant, small to moderate effect sizes (in Cohen’s classification) necessitate relatively large sample sizes (*a priori* power analysis). Based on these considerations, our experiments, especially those testing males for which effect sizes were relatively modest, should have been planned with higher powers to support exercise effects (if true). Therefore, one limitation of our study may reside in the lack of statistical power at detecting such modest effect sizes, a feature that is unfortunately widely diffused in the literature using animal models of neurobehavioral phenomena (Steckler, 2015; McLeod et al. 2015).

We have also found that wheel-running allowed throughout experimentation mitigated the initiation and the expression of psychomotor sensitization to a representative (and hedonistic) dose of cocaine (Brabant et al. 2005), in both female and male mice, and regardless of the period of life in which the exercise period was introduced (either at 28 or 77 days of age). These results reproduce and extend to a sensible post-exercise interval previous findings obtained in our laboratory (Geuzaine and Tirelli, 2014; Lespine and Tirelli, 2015). In contrast, the psychomotor response to the first injection of cocaine (preceding sensitization initiation) was markedly reduced in continuously exercised mice in Experiment 1 (females exercised during adolescence) and Experiment 4 (males exercised during adulthood) but not at all in Experiment 2 (males exercised during adolescence) and Experiment 3 (females exercised during adulthood). In our previous study, in which the main parameters were comparable to those used here, C57BL/6J males with a history of exercise during adolescence did show a clear-cut attenuation of the acute stimulating effects of 8 mg/kg cocaine. These results are not necessarily in contradiction to those of the present Experiment 2 since (1) both sets of results are in the same direction and (2) there is a between-study overlap of the 95% confidence intervals for means and estimates of effect sizes indicating similar plausible values for population parameters. nevertheless, this suggests that in reality, the effect of wheel-running on this phenomenon may be more modest than previously observed. Regarding the responsiveness over the initiation of sensitization (scored as AUC ground) and the expression of the sensitized response to cocaine, we found evidence for preventive properties of continuous wheel-running against cocaine effects, with effect size estimates across experiments varying from moderate to very strong (*d*s ranging from 0.52 to 0.96 for AUC ground and from 0.69 to 1.62 for the sensitization expression). These observations can be considered in line with those reported by Diaz et al. (2013), where male Wistar rats continuously housed with a wheel for at least 5 weeks expressed little or no sensitized psychomotor activity 15 days after 5 once-daily injections of 10 mg/kg cocaine. Smith and Witte (2012) also showed that continuous wheel-running was effective at reducing the psychomotor-activating effects of 3 and 10 mg/kg cocaine in Long-Evans females. Such overall attenuating effect was also observed in C57BL/6J mice housed in large home-cages comprising a running wheel as part of a composite housing environment made of inanimate objects and conspecifics (Bézard et al. 2003; Solinas et al. 2009).

Even if no direct statistical comparisons between ages or genders were carried out the present study (separate experiments), it is clear that wheel-running introduced during adolescence tended to induce larger decreasing effects than when introduced in adulthood, females being more receptive to exercise than males (Cohen’s *d*s of 1.62 vs. 1.02 in females and 0.96 vs. 0.69 in males). This is in substantial accord with the only available study having examined two developmental periods in female rats (Zlebnik et al. 2012). The concurrent access of a running-wheel (available throughout experimentation) and to self-administered cocaine (through lever-pressing) resulted in a decrease in drug seeking among post-weaning adolescents (testing beginning at 23 days of age) but not in adults animals (testing beginning at 90 days of age).

Interestingly, in our males exercised as adolescents, the exercise-induced long-term attenuating effects against cocaine-induced hyperactivity, were significant and rather robust (*d* = 0.53) for the expression of sensitization but non-significant and smaller (*d* = 0.28) for the initiation of sensitization (see Table 1). In females exercised as adolescents, the corresponding effect size for the expression of sensitization was greater (*d* = 1.42) than that found for the initiation of sensitization (*d* = 1.19), both effects being significant. One could hypothesize that these differences might reflect the action of exercise on the processes of sensitization incubation, which takes place during the period preceding the test of expression. Sensitization and its incubation entail alterations of dopaminergic, opioid, serotoninergic and glutamatergic neurotransmissions within the mesolimbic system, a challenge dose being later able to reveal these changes at a behavioral level (Stekette and Kalivas, 2011; Calipari et al. 2015; Leyton and Vezina, 2013; D’Souza, 2015). There is evidence that exercise also acts on this system and possesses rewarding properties, like cocaine and other drugs of abuse (Lett et al. 2000; Belke, 1997; Belke and Wagner, 2005; Greenwood et al. 2011). It is thus tempting to ascribe the exercise-induced protective effects against the cocaine-induced psychomotor sensitization (and its rewarding properties, since we used a rewarding dose of the drug) to a cross-tolerance (from exercise to cocaine effects), at least to some extent. However, given the complexity and multiplicity of the effects of both exercise and cocaine (and the other drugs), one must admit that such an explanatory possibility remains speculative.

Globally, it seems that our females received more beneficial anti-drug effects from exercise than the males. Although we carried out no between-gender statistical comparisons since experiments were realized separately, one can admit that our females exhibited greater levels of wheel-running activity or cocaine responsiveness and stronger interactions between these factors (see Fig. 2). We are aware of no studies addressing gender differences in the long-term effects of exercise on drug responsiveness, and the majority of the available studies assessing drug responsiveness a short time after exercise used only males (Zhou et al. 2016). A greater efficacy of exercise on psychomotor stimulants responsiveness in females, especially in juveniles, has been described for several behavioral measures (for a review see Zhou et al. 2016). This was for example the case for cocaine-primed reinstatement of cocaine self-administration, which was curbed by continuous wheel-running more readily in females (Smith et al. 2012; Zlebnik et al. 2014; however: Smith et al. 2011; Peterson et al. 2014). The greater susceptibility to the cocaine-exercise interaction in our females may result from their greater propensity to exercise, since these tended to wheel-run more than the males (albeit no between-gender statistical comparisons were performed due to the independence of the experiments). This is consistent with many studies previous studies (Lighfoot, 2008; Zhou et al 2016).

## Acknowledgements

The present work was supported by grants from the “Fonds National de la Recherche Scientifique” (FNRS) and the University of Liège “Fonds Spéciaux pour la Recherche” obtained by Ezio TIRELLI. Louis-Ferdinand LESPINE is a research assistant under contract with the FNRS.

